# Using large-scale whole-genome sequence data for single-step genomic predictions in maternal and terminal pig lines

**DOI:** 10.1101/2022.11.11.516229

**Authors:** Sungbong Jang, Roger Ros-Freixedes, John M. Hickey, Ching-Yi Chen, William O. Herring, Ignacy Misztal, Daniela Lourenco

## Abstract

**Background:** Whole-genome sequence (WGS) data harbor causative variants that may not be present in the regular SNP chip data. The objective of this study was to investigate the impact of using preselected variants from WGS for single-step genomic predictions in maternal and terminal pig lines with up to 1.8k sequenced and 104k imputed sequenced animals per line.

**Methods:** Two maternal and four terminal lines were investigated for eight and seven traits, respectively. The number of sequenced animals ranged from 1,365 to 1,491 in maternal lines and 381 to 1,865 in terminal lines. Imputation occurred within each line, and the number of animals imputed to sequence ranged from 66k to 76k in maternal lines and 29k to 104k in terminal lines. Two preselected SNP sets were generated based on genome-wide association study (GWAS). Top40k included the SNP with the lowest p-value in each of 40k genomic windows; ChipPlusSign included significant variants integrated into the regular porcine SNP chip. Single-step genomic predictions with equal or different SNP variances using those SNP sets were compared to the regular porcine SNP chip.

**Results:** In maternal lines, ChipPlusSign, and Top40k showed, on average, 0.62%, and 4.9% increased accuracy compared to the regular porcine SNP chip. The greatest changes were for fertility traits with Top40k, where the initial accuracy based on the SNP chip was low. However, for terminal lines, Top40k resulted in a loss of accuracy of 1% on average. Only ChipPlusSign provided a positive, albeit small, gain (0.85%). Assigning different variances for SNP slightly improved accuracies when using variances obtained from BayesR; however, the increase was inconsistent across the lines and traits.

**Conclusions:** The benefit of using sequence data depends on the line, size of the genotyped population, and how the WGS variants are preselected. When WGS is available on hundreds of thousands of animals, the advantage of sequence data is present but limited in maternal and terminal pig lines.

## Background

Using single nucleotide polymorphisms (SNP) chip data for genomic prediction relies on the linkage disequilibrium (LD) between SNP and causative variants [1]. Because of the initial high cost of SNP genotyping, most of the SNP chips utilized in farm animals are still limited to less than 100k markers, which could restrict the information available for genomic predictions. Whole-genome sequence (WGS) data harbor millions of variants, possibly including causative variants that primarily affect the traits of interest but are not present in regular SNP chips. As sequencing is becoming cheaper, WGS data is becoming available for some agricultural species. Whether this data can help increase the accuracy of genomic predictions beyond that already achieved by SNP chips is still questionable because marginal or no gains were reported by several studies [2-5]. Specifically, in pigs, Zhang et al. [6] showed that the 80k SNP chip outperformed the 650k SNP chip and WGS data for genomic predictions of average daily feed intake and backfat traits. In contrast, Song et al. [7] reported a marginal gain in prediction accuracy when WGS data was used. The absence of benefits reported in those studies could be due to the small number of sequenced animals (maximum of 289 animals), poor imputation accuracy, statistical methods, and sequence SNP that are redundant with the ones in the chip. The largest study about genomic prediction using WGS data in pigs to date, by Ros-Freixedes et al. [8] with almost 400k pigs with imputed WGS data from seven different lines, found small non-robust improvements in prediction accuracy compared to marker arrays for eight common complex traits, and hinted at the need for large datasets and optimized pipelines to take advantage of WGS data.

Imputation is an inevitable step when working with WGS data because sequencing many individuals is still unfeasible. So far, the most efficient approach is to sequence a subset of the animals in a population and impute the sequence to other animals that are genotyped with marker arrays [9]. Not all variants might be causative or in high LD with the causative ones; thus, using the entire WGS data would not benefit genomic predictions [10]. Hence, the preselection of variants helps narrow down the WGS data to only significant ones. Several approaches have been investigated to select significant or causative variants for genomic prediction, such as genome-wide association studies (GWAS) [3], SNP functional annotation [11], and gene expression [12]. Among these approaches, GWAS has been used to preselect WGS variants in pig populations [6-8].

Fragomeni et al. [13] used simulated sequence data to show that once all causative variants are known, and their position and percentage of additive variance explained, prediction accuracy is maximized. Conversely, if only neighboring SNP are identified, the accuracy is inversely proportional to the distance between the causative variants and the neighbor SNP. If only a small proportion of the causative variants were known, the increase in accuracy was also proportional. When a few causative variants were known from a real beef cattle data, Gualdrón-Duarte et al. [14] showed an increase in prediction accuracy for carcass traits of up to seven points when using single-step genomic BLUP (ssGBLUP) with BayesR SNP weights, but no improvements with non-linear weights [2, 15].

Using simulated sequence data, Jang et al. [16] looked at the dimensionality of the genomic information [17] to assess the number of genotyped animals needed to maximize the percentage of discoveries in GWAS. The authors showed that populations with smaller effective size (*Ne* = 20) require more genotyped animals to capture causative variants, whereas for large populations (*Ne* = 200), using the number of genotyped animals equal to that of the largest eigenvalues explaining 98% of the variance of the genomic relationship matrix suffices. Still, only a small proportion of the causative variants can be discovered if those genotyped animals do not have many progeny records.

In pigs, the *Ne* varies from 30 to 50, and the dimensionality of the genomic information or the number of independent chromosome segments segregating in the population ranges from 4k to 6k [18]. Based on Jang et al. [16], using a sample size for GWAS of 7k in a population with *Ne* of 20 allowed identifying causative variants explaining 20% of the additive genetic variance. Moreover, larger sample sizes resulted in larger prediction accuracies with selected variants. Recently, Ros-Freixedes et al. [8], Ros-Freixedes et al. [19] proposed an approach to generate accurate imputed WGS data for hundreds of thousands of pigs across multiple lines and assessed the suitability of WGS variants preselected with GWAS-based methods for genomic prediction using BayesR [20, 21]. However, BayesR considers data solely from genotyped animals. In farm animal populations, only a small fraction of animals is genotyped. In such a situation, single-step methods (i.e., ssGBLUP [22] or ssSNP-BLUP [23]) are advantageous because they also incorporate information on non-genotyped individuals. Additionally, most breeding programs currently use single-step methods [24, 25]. Therefore, the present study investigates the impact of using preselected variants from WGS data for genomic prediction with ssGBLUP in maternal and terminal pig lines, with up to 1.8k sequenced and 104k imputed sequenced animals per line. We explored different ways to preselect variants and the changes in accuracy when using ssGBLUP and weighted ssGBLUP (WssGBLUP) with BayesR SNP variances as weights.

## Methods

### Data

Datasets provided by the Pig Improvement Company (PIC; Hendersonville, TN) comprised two maternal lines (ML1, ML2) and four terminal lines (TL1, TL2, TL3, and TL4) with diverse genetic backgrounds, which breeds of origin were Landrace and Large White for maternal lines, and Duroc, Hampshire, and Large White for terminal lines. For maternal lines, we investigated average daily feed intake (ADFI), average daily gain (ADG), backfat thickness (BF), loin depth (LDP), total number of piglets born (TNB), number of stillborn (NSB), return to oestrus seven days after weaning (RET), and litter weaning weight (WWT). For terminal lines, we investigated ADFI in purebreds, but ADG, BF, and LDP in both purebreds and crossbreds (ADGX, BFX, and LDPX). Some traits were jointly analyzed in multi-trait models for genomic prediction. In maternal lines, two-trait models were considered for ADG and ADFI (ADFI model), ADG and BF (GROWTH model), ADG and LDP (LOIN model), TNB and NSB (REPROD model), whereas single-trait models were used for RET (RET model) and WWT (WWT model). In terminal lines, the same ADFI model as for the maternal lines was applied, but four-trait models were used for the GROWTH (ADG, BF, ADGX, and BFX) and LOIN (ADG, LDP, ADGX, and LDPX) models. The total number of animals in the pedigree and records for each trait are in Table 1. Pigs were genotyped with the GGP-Porcine HD BeadChip (GeneSeek, Lincoln, NE). We filtered out the monomorphic SNP as well as SNP with a call rate lower than 0.90, minor allele frequency lower than 0.01, and the difference between observed and expected genotype frequencies greater than 0.15. Individuals with more than 10% missing genotypes were also removed. Table 2 depicts the number of genotyped animals and SNP per line after quality control.

**Table 1.**
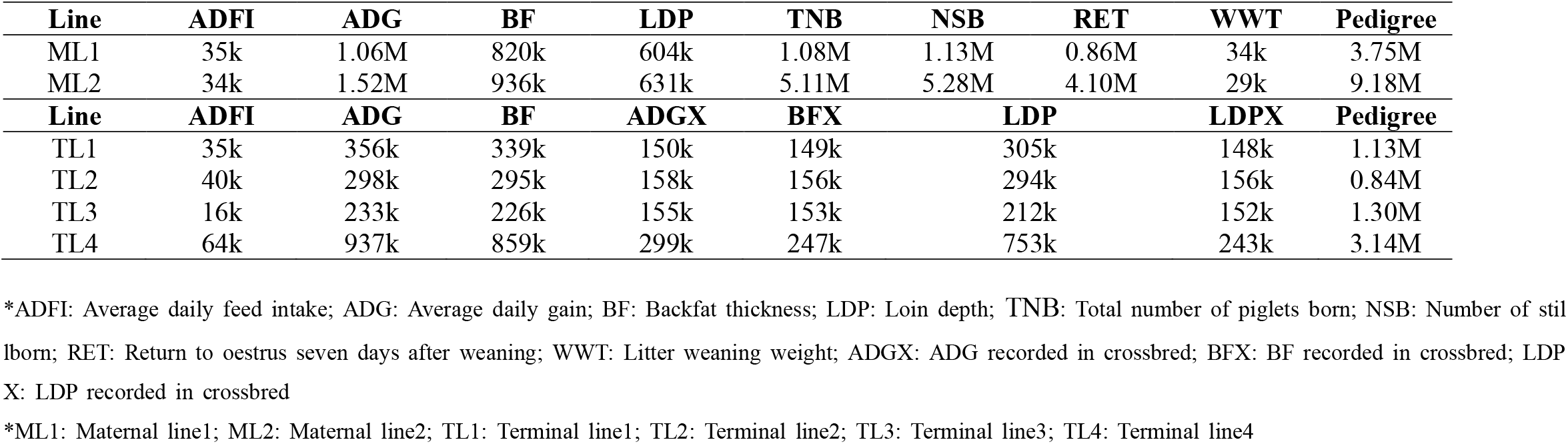
Number of records and animals in the pedigree.

| Line | ADFI | ADG | BF | LDP | TNB | NSB | RET | WWT | Pedigree |
| --- | --- | --- | --- | --- | --- | --- | --- | --- | --- |
| ML1 | 35k | 1.06M | 820k | 604k | 1.08M | 1.13M | 0.86M | 34k | 3.75M |
| ML2 | 34k | 1.52M | 936k | 631k | 5.11M | 5.28M | 4.10M | 29k | 9.18M |
| Line | ADFI | ADG | BF | ADGX | BFX | LDP | LDPX | Pedigree |  |
| TL1 | 35k | 356k | 339k | 150k | 149k | 305k | 148k | 1.13M |  |
| TL2 | 40k | 298k | 295k | 158k | 156k | 294k | 156k | 0.84M |  |
| TL3 | 16k | 233k | 226k | 155k | 153k | 212k | 152k | 1.30M |  |
| TL4 | 64k | 937k | 859k | 299k | 247k | 753k | 243k | 3.14M |  |

\*ADFI: Average daily feed intake; ADG: Average daily gain; BF: Backfat thickness; LDP: Loin depth; TNB: Total number of piglets born; NSB: Number of stillborn; RET: Return to oestrus seven days after weaning; WWT: Litter weaning weight; ADGX: ADG recorded in crossbred; BFX: BF recorded in crossbred; LDPX: LDP recorded in crossbred
\*ML1: Maternal line1; ML2: Maternal line2; TL1: Terminal line1; TL2: Terminal line2; TL3: Terminal line3; TL4: Terminal line4

**Table 2.** Number of genotyped individuals, SNP, sequenced, and imputed sequenced animals in six lines.

| **Line** | **Number of genotyped individuals** | **Number of SNP (Chip)** | **Number of sequenced individuals** | **Number of imputed sequenced individuals** |
| --- | --- | --- | --- | --- |
| ML1 | 76,227 | 40,592 | 1,366 | 76,230 |
| ML2 | 66,608 | 42,746 | 1,491 | 66,608 |
| TL1 | 60,467 | 35,786 | 731 | 60,474 |
| TL2 | 41,572 | 40,311 | 760 | 41,573 |
| TL3 | 29,328 | 39,999 | 381 | 29,330 |
| TL4 | 104,644 | 43,032 | 1,856 | 104,661 |

\*ML1: Maternal line1; ML2: Maternal line2; TL1: Terminal line1; TL2: Terminal line2; TL3: Terminal line3; TL4: Terminal line4

### Whole-genome sequencing and imputation

The WGS data used in this study were generated by Ros-Freixedes et al. [8], Ros-Freixedes et al. [19]. In summary, a low-coverage sequencing strategy was followed by joint calling, phasing, and imputation of the WGS genotypes using the ‘hybrid peeling’ method implemented in AlphaPeel [26]. The number of sequenced and imputed sequenced individuals for each line is provided in Table 2. The ‘hybrid peeling’ method used all the marker arrays (GGP-Porcine LD and HD) and WGS data available across complete multi-generational pedigrees. Imputation was carried out separately for each line. Individuals were predicted to have low imputation accuracy if they or their grandparents were not genotyped with a marker array or if they were less connected (based on pedigree relationships) to the rest of the population. Individuals with low predicted imputation accuracy were excluded, as described by Ros-Freixedes et al. [19]. A total of 76,230 (ML1), 66,608 (ML2), 60,474 (TL1), 41,573 (TL2), 29,330 (TL3), and 104,661 (TL4) imputed individuals remained in each line after quality control (Table 2). These individuals were predicted to have an average dosage correlation of 0.97 (median: 0.98) based on the imputation accuracy of 284 pigs that had both WGS (high coverage) and marker array data. All SNP with a minor allele frequency lower than 0.023 were removed since their estimated dosage correlations were lower than 0.90 [19]. Genotypes were called after imputation even for the individuals that were directly sequenced, to account for information from the relatives in genotype calling and reduce uncertainty from low coverage.

### Training and test sets

Before the GWAS, all animals with WGS data were separated into training and test, which were defined as in Ros-Freixedes et al. [8]. Test sets were generated by extracting entire litters from the last generation of the pedigree. The training sets were created by establishing a threshold on the pedigree relationship coefficients between training and test sets. We removed individuals with a relationship coefficient equal to or greater than 0.5 to test animals to resemble the selection candidate evaluation done by pig breeding companies. The same training set was used for GWAS and genomic predictions in each line. Previous studies reported a reduction in prediction accuracy and bias of genomic estimated breeding value (GEBV) when using the same dataset for GWAS and genomic prediction [27, 28]. However, Ros-Freixedes et al. [8], using the same data as in our study, revealed no systematic changes after splitting the training set into two exclusive subsets, one for each step of the analysis.

### Pre-selected SNP panels

Two different pre-selected SNP panels were created from WGS for the genomic prediction as described in Ros-Freixedes et al. [8]: (1) Top40k and (2) ChipPlusSign. Top40k were the variants with the lowest p-value (not necessarily below the significance threshold) in each consecutive non-overlapping 55-kb window along the genome, based on GWAS. ChipPlusSign combined the GGP-Porcine HD SNP and significant variants (p ≤ 10^−6^) from the GWAS in a way that when a 55-kb window contained more than one significant variant, only that with the lowest p-value was selected. Genomic predictions with the two sets were compared against the GGP-Porcine HD chip (Chip). For scenarios that used multi-trait models, the preselected variants for each trait were combined for the traits included in each model. For example, those pre-selected variants for each ADFI and ADG were combined for the ADFI model and used for genomic prediction. As WGS information was available only on purebred animals, no variants were selected for ADGX, BFX, and LDPX. All the combinations of selected variants in each model are described in Additional file 1: Table S1. After constructing all the pre-selected SNP panels, quality control was done to remove SNP with a difference between observed and expected genotype frequencies greater than 0.15 and to exclude individuals with parent-progeny Mendelian conflicts. Additional file 1: Table S2 and S3 depict the number of animals and SNP for all pre-selected SNP panels after quality control in the maternal and terminal lines.

Because the number of animals available for each preselected SNP panel (Chip, Top40k, and ChipPlusSign) was different after quality control, the training and validation populations contained only the common animals that passed quality control for all the SNP panels. This step guaranteed a fair comparison of genomic prediction (Table 3).

**Table 3.** Number of animals with genomic information that were retained after quality control and used in the analyses with all the SNP panels.

| Line | ADFI | GROWTH | LOIN | REPROD | RET | WWT |
| --- | --- | --- | --- | --- | --- | --- |
| ML1 | 74,148 | 74,153 | 74,152 | 73,919 | 73,891 | 74,058 |
| ML2 | 64,654 | 64,655 | 64,659 | 64,599 | 64,653 | 63,456 |
| TL1 | 56,423 | 56,424 | 56,422 | - | - | - |
| TL2 | 38,477 | 38,475 | 38,477 | - | - | - |
| TL3 | 27,671 | 27,671 | 27,671 | - | - | - |
| TL4 | 102,586 | 102,590 | 102,588 | - | - | - |

\*In ML1 and ML2; ADFI: two-trait ADFI model (ADG and ADFI); GROWTH: two-trait GROWTH model (ADG and BF); LOIN: two-trait LOIN model (ADG and LDP); REPROD: two-trait REPROD model (TNB and NSB); RET: single-trait RET model (RET); WWT: single-trait WWT model (WWT)
\*In TL1, TL2, TL3, and TL4; ADFI: two-trait GROWTH model (ADG and ADFI); GROWTH: four-trait GROWTH model (ADG, BF, ADGX, and BFX); LOIN: four-trait LOIN model (ADG, LDP, ADGX, and LDPX)

### Genomic prediction

Single-trait, two-trait, or four-trait models were used for genomic prediction, depending on the traits. Herein, only the four-trait GROWTH model (ADG, BF, ADGX, and BFX) of terminal lines is described:

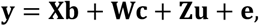

where **y** is the vector of phenotypes; **X** is an incidence matrix for fixed effects (contemporary group as a cross-classified effect for all traits, off-test weight and carcass weight as a covariate only for BF and BFX, respectively) contained in **b**; **W** is an incidence matrix for the random, diagonal litter effect contained in **c** (**c** ∼ MVN(0, **I⨂L**_0_)); **Z** is an incidence matrix for the random additive genetic effect contained in **u** (**u** ∼ MVN(0, **H⨂Σ**_0_)); and **e** (**e** ∼ MVN(0, **I⨂R**_0_)) is a vector of residual effects. Matrices **L**_0_, **Σ**_0_, and **R**_0_ are as follows:

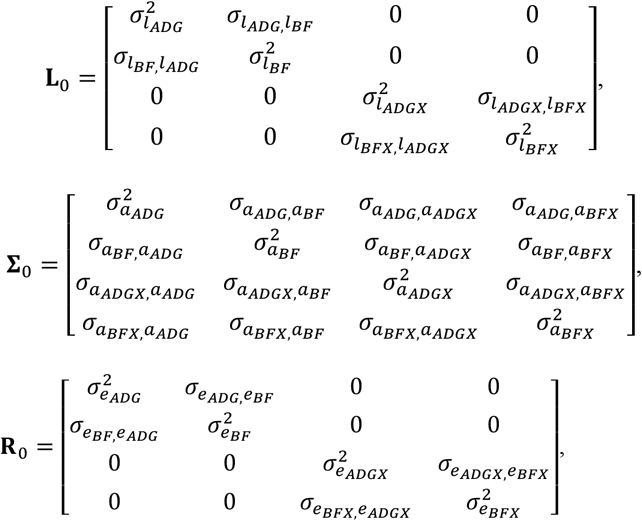

where 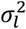 is the litter variance, 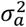 is the additive genetic variance, and 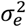 is the residual variance. **I** is an identity matrix and **H** is the realized relationship matrix that combines pedigree and genomic relationships in ssGBLUP. The genomic prediction was performed with both ssGBLUP and WssGBLUP using the BLUPF90 family of programs [29], which used the inverse of **H** (**H**^−1^) as follows [30]:

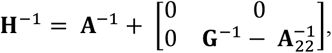

where **G**^−1^ is the inverse of the genomic relationship matrix, **A**^−1^ and 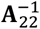 are the inverses of the pedigree relationship matrices for all and genotyped individuals, respectively. The **G** was created using the first method of VanRaden [31]:

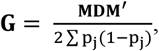

where **M** is a matrix of genotypes centered for current allele frequencies, p_j_ is the minor allele frequency of SNP j, and **D** is the diagonal matrix of SNP weights. All the SNP were presumed to have homogeneous weights in ssGBLUP, meaning that **D** is an identity matrix (**I**). To ensure compatibility between **G** and **A**_22_ and circumvent singularity issues, **G** was tuned and then blended with 5% of **A**_22_.

The algorithm for proven and young (APY) was applied to obtain **G**^−**1**^ while avoiding the direct inversion of **G** [32] for the lines with more than 50k genotyped animals, i.e., ML1, ML2, TL1, and TL4. Lines TL2 and TL3 used direct inversion of **G**. To ensure reliable estimation of GEBV, the number of core animals corresponded to the number of largest eigenvalues explaining 98% of the total variation in **G** assessed with regular chip data [17]. Therefore, the number of core animals in each line was: 4,200, 5,400, 3,400, and 5,500 for ML1, ML2, TL1, and TL4, respectively.

For WssGBLUP, we calculated SNP variances from BayesR [20] and assigned those variances as weights for SNP in an iterative way. Each iteration stored individual SNP variances, and posterior SNP variance was calculated as the average variance across all the iterations. Afterward, the weights were re-scaled to make the trace of **D** equal to the number of SNP. More details about BayesR weighting are described in Gualdrón-Duarte et al. [14]. This approach was only applied to the four largest lines, ML1, ML2, TL1, and TL4 for growth-related traits (ADFI, ADG, BF, and LDP) with the Top40k and ChipPlusSign data.

### Validation

The accuracy of genomic prediction was calculated by correlating GEBV with deregressed EBV (dEBV) [33] for the animals in the test sets. Inflation or deflation levels were assessed as the b_1_ of the regression of dEBV on GEBV. The b_1_ values lower than 1 indicated inflation of GEBV and greater than 1 indicated deflation. All animals with no dEBV were removed from the test sets. The number of test animals for each trait in all lines is summarized in Additional file 1: Table S4.

## Results

### Genomic prediction accuracy of maternal lines using ssGBLUP

Fig. 1 shows the changes in prediction accuracy (%) of ChipPlusSign, and Top40k compared to Chip for the two maternal lines. All results regarding prediction accuracy and changes respective to Chip (%) are summarized in Additional file 1: Tables S5 and S6, respectively. For many traits, ChipPlusSign and Top40k showed greater accuracy than Chip (Fig. 1). Using ChipPlusSign instead of Chip resulted in a maximum gain of 1.61% for ADG in ML1 and 1.49% for ADFI in ML2, and the greatest decrease of -0.26% and -0.74% for WWT in ML1 and ML2, respectively. The mean across all eight traits decreased as the number of genotyped animals decreased from ML1 (76,227) to ML2 (66.608), although the percentage of gain was very small (0.75% to 0.49%) compared to Chip. Accuracy gains by using Top40k were greater than the ones from ChipPlusSign. The average gain using Top40k for ML1 and ML2 was 5.54% and 4.34%, respectively, following the decreasing number of animals with WGS data. The largest gains in each line with Top40k were 34.77% and 22.87% for RET in ML1 and ML2, whereas the largest reduction was -5.19% and -4.25% for WWT in ML1 and ADFI in ML2, respectively. Therefore, pre-selection of variants from GWAS (ChipPlusSign and Top40k) could improve prediction accuracy for most traits in maternal lines, although those gains were small to modest.

**Figure 1.**
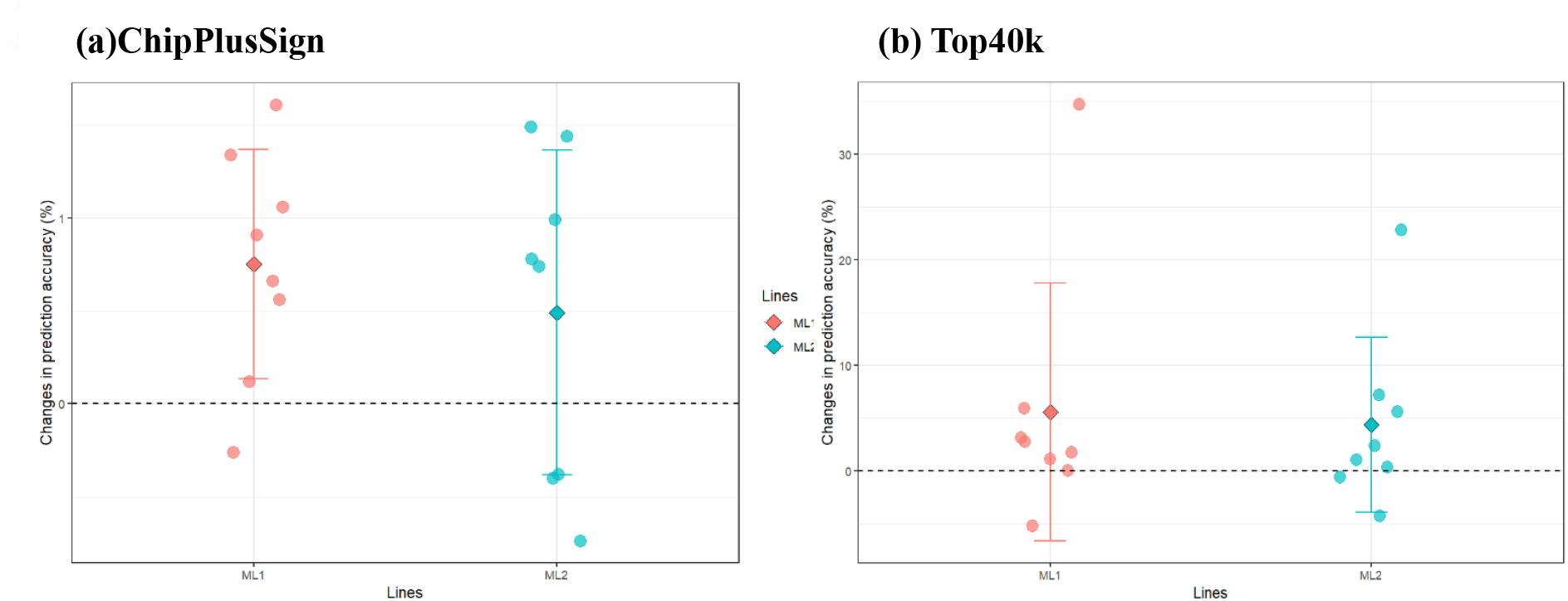
Accuracy changes (%) of ChipPlusSign and Top40k compared to Chip in maternal lines. *Each circle represents accuracy changes in each trait, whereas diamonds indicate the mean accuracy change across all traits in each line *(a) ChipPlusSign: genomic prediction using ChipPlusSign preselected genotype panel *(b) Top40k: genomic prediction using Top40k preselected genotype panel

### Genomic prediction accuracy of terminal lines using ssGBLUP

Changes in prediction accuracy (%) of ChipPlusSign, and Top40k compared to Chip for four terminal lines are described in Fig. 2. All results regarding prediction accuracy and changes respective to Chip (%) are summarized in Additional file 1: Tables S7 and S8. ChipPlusSign reported a consistent gain in accuracy among all traits and lines, except for LDPX in TL2 (−5.26%) and LDPX in TL3 (−0.45%). As the results of ChipPlusSign in maternal lines, the result in terminal lines also showed a decreasing pattern as the number of genotyped animals decreased, in the following order: TL4 (104,644), TL1 (60,467), TL2 (41,572), and TL3 (29,328). The average gain for all seven traits was 0.95%, 0.56%, 0.37%, and 1.50% (TL1 to TL4) and the maximum gain was 1.32% (TL1-BF), 2.27% (TL2-ADGX), 0.94% (TL3-ADFI), and 2.61% (TL4-ADGX). Contrary to ChipPlusSign results, Top40k reported inconsistent results among the traits and lines. Although TL3 and TL4 showed accuracy gains for most traits, except ADFI in TL4 (−0.49%), TL1 and TL2 reported a reduction in accuracy for many traits (six traits for TL1 and four traits for TL2). On average, TL3 showed the second greatest accuracy gain for all seven traits (2.35%), with the smallest number of genotyped animals among all terminal lines. In contrast, TL1 reported the most considerable reduction in accuracy, even though this line is the second largest genotyped line. TL1, TL2, and TL4 reported -6.28%, -3.32%, and 3.30% accuracy changes, meaning that the number of genotyped animals did not affect the gain with Top40k for terminal lines. However, the largest genotyped line (TL4) showed the most notable average gain (3.30%). The maximum gains in each line were 1.55% (BFX). 16.16% (ADGX), 3.74% (LDP), and 7.91% (ADGX) for TL1, TL2, TL3, and TL4, respectively. For both ChipPlusSign and Top40k, TL2 showed the greatest standard deviation among the traits, 2.67 and 18.37, respectively, meaning that accuracy changes highly depend on the traits in TL2. In general, the use of ChipPlusSign reported improved accuracy for most of the traits in the terminal lines. However, those gains were limited (maximum 2.61% for ADGX in TL4). Results of Top40k showed decreased accuracy in most of the traits for TL1 and TL2, on the contrary, increased accuracy was outlined in TL3 and TL4 for almost all traits.

**Figure 2.**
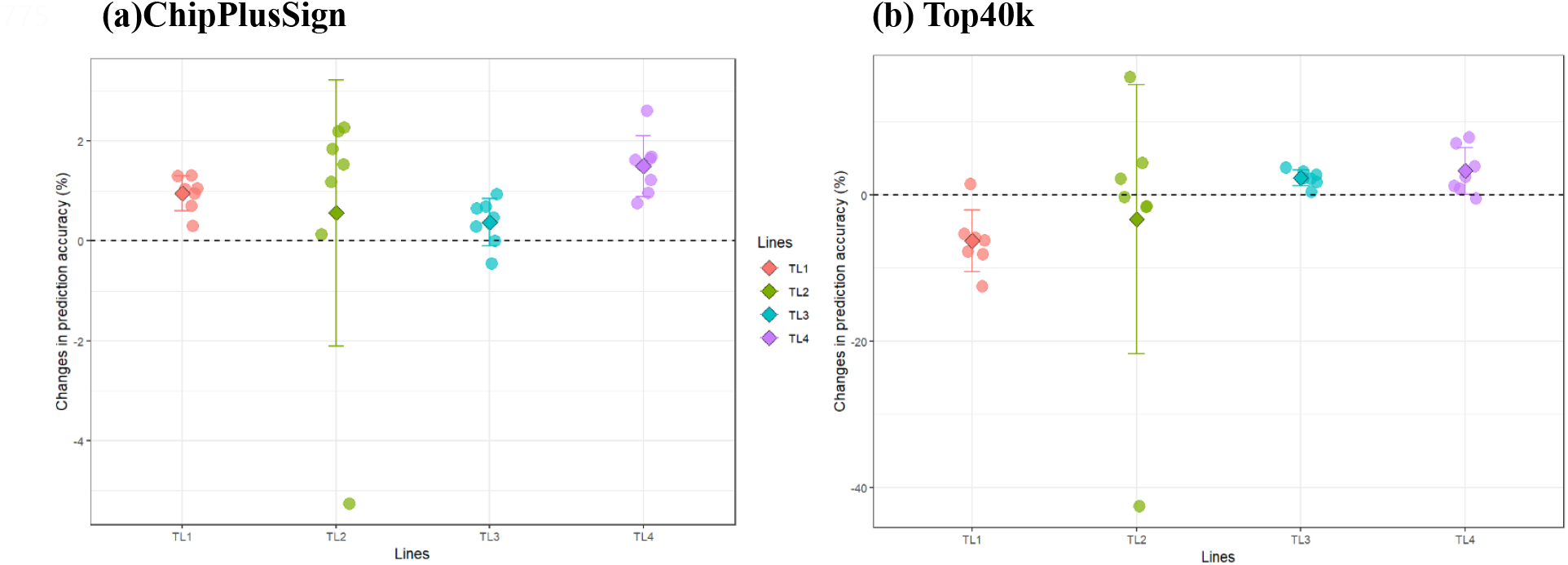
Accuracy changes (%) of ChipPlusSign and Top40k compared to Chip in terminal lines. *Each circle represents accuracy changes in each trait, whereas diamonds indicate the mean accuracy change across all traits in each line *(a) ChipPlusSign: genomic prediction using ChipPlusSign preselected genotype panel *(b) Top40k: genomic prediction using Top40k preselected genotype panel

### Inflation/deflation of GEBV

Fig. 3 describes the b_1_ values for all genotyped scenarios for maternal (a) and terminal lines (b). All the values in each genotype scenario were averaged across all traits in each line. The specific results for b_1_ of all maternal lines and terminal lines were summarized in Additional file 1: Tables S9 and S10, respectively. Fig. 3-(a) described the results of maternal lines. When the number of genotyped animals decreased (ML1 to ML2), b_1_ got closer to 1.0 (0.68 in ML1 and 0.81 in ML2). More specifically, all genotype panels had smaller inflation of GEBV from ML1 to ML2. The results of Chip, ChipPlusSign, and Top40k were similar within the lines.

**Figure 3.**
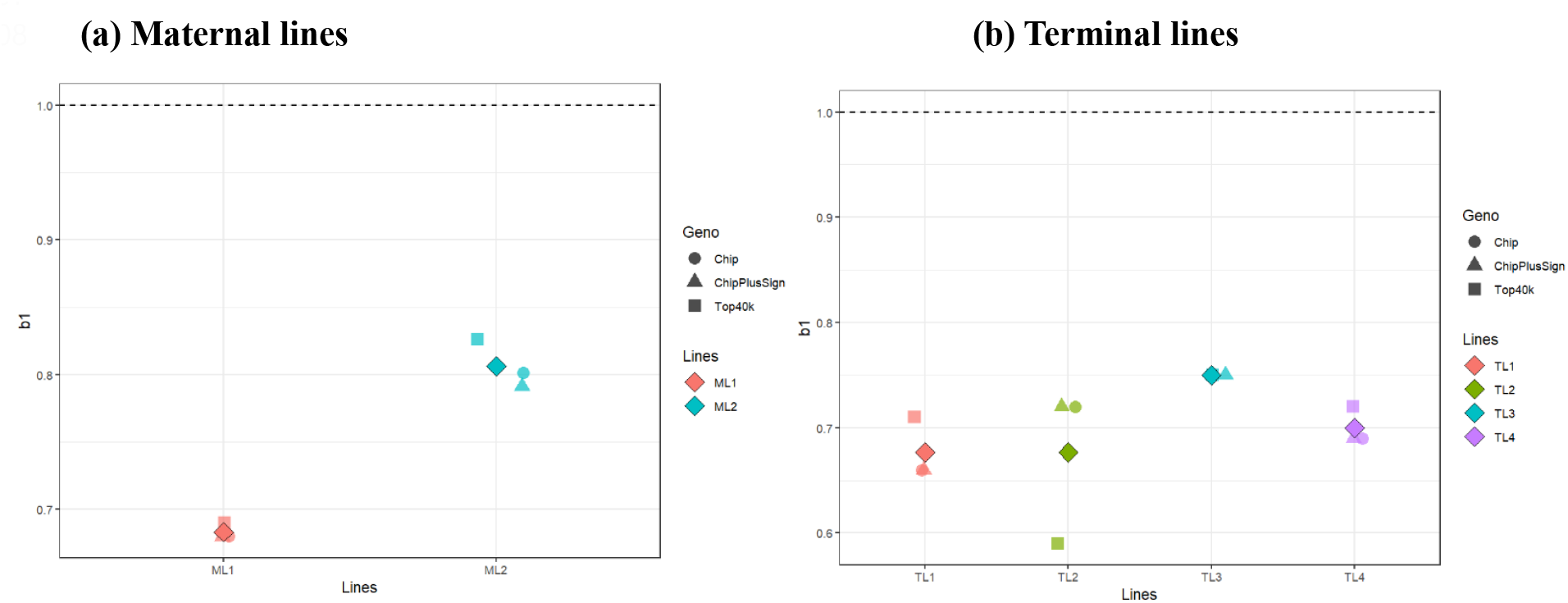
b_1_ values for all the genotype scenarios in both maternal and terminal lines. *Diamond shape indicated the overall mean of b_1_ values for all traits and genotype panels in each line

The results of terminal lines are in Fig. 3-(b). Compared to the results of maternal lines, inconsistent patterns were observed in terminal lines among the traits and lines. Overall, all the terminal lines showed inflated GEBVs for all traits and genotype panels. On average, TL3 reported the best result (0.75), followed by TL4, TL1, and TL2 (0.70, 0.68, and 0.68, respectively). The Chip, ChipPlusSign, and Top40k yielded similar results in TL3 and TL4. In TL1 and TL2, however, Top40k showed either less inflation (+0.05) or more inflation (−0.13) compared to Chip and ChipPlusSign.

### Genomic prediction using WssGBLUP

WssGBLUP using BayesR weights was only applied to the four largest lines (ML1, ML2, TL1, and TL4) for growth-related traits (ADFI, ADG, BF, and LDP) with Top40k and ChipPlusSign. Results of prediction accuracy and b_1_ are in Table 4 and Additional file 1: Table S11, respectively. For ML1, no gain was observed with WssGBLUP. In ML2, using Top40k and ChipPlusSign with WssGBLUP showed a 0.02 accuracy gain in BF compared to ssGBLUP, whereas -0.09 reduction in LDP. TL1 results also showed no improvement in accuracy. The greatest gains (∼0.06) were outlined in the results of TL4 when using WssGBLUP compared to ssGBLUP. No gain was observed for ADFI. However, 0.06, 0.03, and 0.04 accuracy gains were observed for ADG, BF, and LDP when Top40k weighted was used instead of Top40k. Similarly, 0.03, 0.02, and 0.04 increases in accuracy were reported with ChipPlusSign and BayesR weights for ADG, BF, and LDP. Overall, WssGBLUP showed similar b_1_ values to the regular ssGBLUP except for a few scenarios (results not shown).

**Table 4.** Prediction accuracy of WssGBLUP compared to ssGBLUP.

| **Line** | **Description** | **ADFI** | **ADG** | **BF** | **LDP** |
| --- | --- | --- | --- | --- | --- |
| ML1 | Top40k | 0.37 | 0.49 | 0.51 | 0.53 |
|  | Top40k weighted | 0.37 | 0.49 | 0.51 | 0.53 |
|  | ChipPlusSign | 0.37 | 0.47 | 0.51 | 0.52 |
|  | ChipPlusSign weighted | 0.37 | 0.47 | 0.51 | 0.51 |
| ML2 | Top40k | 0.36 | 0.61 | 0.64 | 0.62 |
|  | Top40k weighted | 0.35 | 0.62 | 0.66 | 0.53 |
|  | ChipPlusSign | 0.37 | 0.61 | 0.63 | 0.62 |
|  | ChipPlusSign weighted | 0.37 | 0.62 | 0.65 | 0.53 |
| TL1 | Top40k | 0.34 | 0.45 | 0.59 | 0.56 |
|  | Top40k weighted | 0.34 | 0.45 | 0.59 | 0.55 |
|  | ChipPlusSign | 0.36 | 0.49 | 0.61 | 0.60 |
|  | ChipPlusSign weighted | 0.36 | 0.50 | 0.60 | 0.60 |
| TL4 | Top40k | 0.39 | 0.51 | 0.60 | 0.59 |
|  | Top40k weighted | 0.39 | 0.57 | 0.63 | 0.63 |
|  | ChipPlusSign | 0.40 | 0.51 | 0.60 | 0.57 |
|  | ChipPlusSign weighted | 0.40 | 0.54 | 0.62 | 0.61 |

\*ML1: Maternal line1; ML2: Maternal line2; TL1: Terminal line1; TL4: Terminal line4
\*ADFI: Average daily feed intake; ADG: Average daily gain; BF: Backfat thickness; LDP: Loin depth
\*Top40k: Top40k preselected genotype panel; Top40k weighted: Top40k using BayesR weighting
\*ChipPlusSign: ChipPlusSign preselected genotype panel; ChipPlusSign weighted: ChipPlusSign using BayesR weighting

## Discussion

The current study investigated the impact of using large-scale WGS data for genomic prediction through ssGBLUP and WssGBLUP in maternal and terminal pig lines. This is the first study applying ssGBLUP to large-scale WGS pig datasets, with up to 1.8k sequenced and 104k imputed sequenced animals per line. Two sets of preselected WGS variants were created to compare genomic predictions with WGS and the regular SNP chip. Our results showed that preselected variants could outperform the regular SNP chip for genomic prediction, although not consistently across the lines and traits and with relatively limited gains, as it was also observed using a BayesR [8]. In addition, we observed the potential to improve prediction accuracy through WssGBLUP using posterior variance from the BayesR as SNP weight, especially for the largest genotyped populations. Our results suggest effective scenarios to construct preselected variant sets depending on traits and population sizes for maternal and terminal pig lines. In the discussion section, we will address the three topics: (1) Impact of the method for preselecting WGS variants on genomic prediction (2) Using WGS data for genomic prediction in pigs (3) Comparison of weighted and non-weighted ssGBLUP.

### Impact of the method for preselecting WGS variants on genomic prediction

Theoretically, using WGS data can improve genomic predictions because they cover the entire genome, assuming that causative variants are likely presented in the data. Therefore, genomic prediction does not rely on LD between SNP and causative variants but directly uses causative variants [34]. A common assumption is that all the variants can explain a large proportion of genetic variance in the WGS data. However, several studies reported that using all variants in WGS data did not improve prediction accuracies [6, 7, 10]. A plausible reason for that would be the use of redundant SNP. As the WGS data have millions of SNP across the entire genome, adjacent SNP likely have a strong LD with causative variants or other SNP in certain genome blocks, indicating that most SNP are correlated, providing the same information. Therefore, fitting all the WGS data in the prediction model could lead to biased GEBV. To avoid bias, many studies have investigated the preselection of significant variants for genomic prediction [2-4].

Thus, in the current study, two different preselected genotype panels were designed and compared to the regular chip data. Those panels were constructed following different assumptions as in Ros-Freixedes et al. [8], Ros-Freixedes et al. [35]. For ChipPlusSign, significant variants (p≤10^−6^) based on GWAS were added to the regular SNP chip with an expectation of better prediction accuracy if the significant SNP had large effects or were causative and not presented in Chip. Incorporating preselected, significant SNP into the regular SNP chip has been investigated in many studies with WGS data [3, 13, 36]. Top40k was created to mimic the number of SNP in the regular medium-density SNP chips used for routine genomic evaluation in many farm animals (e.g., pigs, cattle, and chickens). As most of the regular SNP chips contain evenly spaced SNP, Top40k also consisted of 40k SNP selected from each consecutive non-overlapping 55-kb window of WGS with the lowest p-value (i.e., from GWAS) in each window. Therefore, gains in prediction accuracy were expected if those preselected 40k SNP from WGS data were more informative and explained a more notable proportion of genetic variation than the SNP in the regular chip. In this study, Top40k sets in the multi-trait models (ADFI, GROWTH, and LOIN) were combined, generating around 80k SNP (Additional file 1: Table S2 and Table S3), which was differently handled by Ros-Freixedes et al. [8] who used the same dataset but only with single-trait models.

Among the preselected genotype sets, ChipPlusSign showed small to moderate accuracy gain for many traits in maternal and terminal lines. This panel also showed the most consistent results across lines and traits, with accuracy gains in most of them; however, within a limited range (from 0.12% to 2.61%). ChipPlusSign also showed greater robustness than Top40k when the genomic prediction was performed using BayesR [8]. Several studies have been conducted to investigate genomic prediction by adding selected variants to regular chip data through either real or simulated datasets [2, 3, 16, 36]. In US Holstein, VanRaden et al. [3] investigated the reliability of GEBV for 33 traits when preselected SNP (N = 16k) from WGS were added to a 60k SNP chip. They reported up to a 4.8 percentage point increase in reliability (15.35%) with an average of 2.7 extra points (9.15%) compared to the reliability obtained from a 60k SNP chip. However, when Fragomeni et al. [2] investigated the performance of ssGBLUP using the same preselected variants set created by VanRaden et al. [3], almost no gain in reliability (0.92%) was observed, although reliabilities were greater in Fragomeni et al. [2]. One major difference between those two studies was the method of genomic prediction, multistep in VanRaden et al. [3] and ssGBLUP in Fragomeni et al. [2]. When ssGBLUP was used, it combined all the information from genotyped and non-genotyped animals, allowing the inclusion of a massive amount of data. In such a scenario, gains in reliability are less likely if the selected variants are redundant, not truly causative, or have a small effect on the traits of interest. Our results agree with the ones from Fragomeni et al. [2], especially with ChipPlusSign. In a simulated study, Jang et al. [16] investigated the dimensionality of genomic information for variant selection and genomic prediction with sequence data. Their results showed that populations with small *Ne* obtained a maximum accuracy gain of 0.86% to 1.98% when either significant variants or hundreds of variants with high effect sizes preselected from GWAS were added to a 50k SNP chip. The small *Ne* scenario they simulated was 20, close to the *Ne* in pig populations (32∼48) [18].

In our study, the results of Top40k highly depended on the traits and lines. Top40k showed the greatest gain for RET across all maternal lines (22.87% to 34.77%), but relatively marginal gains or reductions for other traits. In the terminal lines, results of Top40k fluctuated more among lines, with increased or decreased accuracies. The possible reason for the large improvement observed for reproduction or fertility traits in maternal lines might be the nature of the traits and the lack of informative SNP in the regular SNP chip. The lack of informative SNP for fertility traits led to a recent change in the SNP chip for beef cattle evaluations (https://www.angus.org/AGI/global/AngusGS.pdf).

Heritabilities for RET were relatively lower than other traits. Consequently, this trait had the lowest prediction accuracies (0.14 for ML1 and 0.20 for ML2 with Chip) among other traits in maternal lines. Thus, there would be greater room for improvement in genomic predictions using preselected genotype data if the SNP in Chip could not explain a large proportion of the genetic variance. Therefore, we speculated that there would be more informative SNP in Top40k for RET, which were not included in the regular SNP chip. Likewise, small-scale differences in accuracy observed between Chip and Top40k were possibly due to variants capturing similar proportions of genetic variance and having similar LD patterns across the genome. In terminal lines, the maximum gain was observed for ADGX (16.16% in TL2), recorded in crossbred animals. The ADGX was investigated in the GROWTH model along with three correlated traits (ADG, BF, and BFX), and the Top40k was created based on GWAS for ADG and BF, individually, and then each Top40k was combined for genomic prediction. Thus, this result showed the potential for prediction improvement for the traits recorded in crossbred animals if many phenotypes and WGS animals are available although those traits were not directly used for preselection of variants.

Those marginal gains for most traits with ChipPlusSign and Top40k raised a question about the amount of information that has been used for preselecting the variants and performing genomic predictions. Examining the dimensionality of the genomic information can help assess the efficient number of genotyped animals needed to maximize the percentage of discoveries in GWAS and prediction accuracy gains [16]. According to Jang et al. [16], in populations with a larger effective size (*Ne* = 200), using the number of genotyped animals equal to the number of largest eigenvalues explaining 98% of the variance of **G** sufficed to capture the most informative variants, although only a tiny proportion of the causative variants was discovered for highly polygenic traits. However, their study showed that populations with a smaller effective size (*Ne* = 20) required much more data to capture causative variants. For example, when 30k genotyped animals were used in GWAS for highly polygenic traits, only three causative variants were identified, explaining 3.9% of genetic variation. In addition, incorporating preselected variants to regular chip data reported a nearly 2% maximum gain in accuracy for the scenarios with *Ne* = 20. In the current study, the number of WGS animals used for GWAS was 29k to 104k, which is the largest WGS data in pigs by far. However, the fine-mapping of causative variants was still challenging and the benefits for genomic predictions were limited [8]. Since pig populations have small *Ne* and most of the traits are highly polygenic, to capture the most informative variants, a very large number of WGS animals having lots of progeny records would be required [16].

In an initial batch of analyses (results not shown), we used only significant variants (TopSign) for genomic predictions, which showed no benefits compared to Chip. In fact, accuracies were reduced for most of the traits and lines. The number of variants in TopSign ranged from 6 to 1,705 depending on the lines and traits. Fragomeni et al. [13] outlined that the maximum accuracy of GEBV could be obtained if the true causative variants were identified with their exact substitution effects, position in the genome, and genetic variance explained by each variant assigned as weight. Therefore, our results revealed that the variants in TopSign might not be the true causative variants, so the use of those variants underperformed regular SNP chip information.

### Using WGS data for genomic prediction in pigs

The cost of sequencing is getting cheaper, so using sequence data for genomic prediction of farm animals (e.g., sheep, beef cattle, dairy cattle, and pigs) has been more approachable than in the past. Several studies have been carried out and reported marginal or no benefits of using WGS on genomic prediction in sheep, beef, and dairy cattle [2, 3, 11, 12, 36]. Compared to other farm animals, there are scarce studies on the use of WGS data for genomic prediction in other populations of pigs, in which only small-scale datasets were used [6, 7]. The number of imputed sequenced pigs in those studies was less than 7k. As the number of variants in WGS increases, more samples are required to resolve the well-known issue of ‘*N << p*’, where *N* is the sample size and *p* is the number of variants. If the sample size is not sufficient, estimation of SNP effects and identification of causative SNP could be troublesome, especially for populations with small *Ne* and highly polygenic traits.

In the current study, the number of sequenced animals was around 380 to 1.8k across lines, which represented nearly 2% of the population in each line [8, 35]. However, depending on the line, the WGS information was imputed for 29k to 104k animals. Applying large-scale WGS data to preselect the variants through GWAS and using those variants for genomic prediction in our study and Ros-Freixedes et al. [8] showed limited improvement as found in Zhang et al. [6], Song et al. [7], in which only a small number of animals had WGS. Theoretically, increasing the sample size enhanced the power to detect causative variants and improved genomic predictions [34, 37]. However, the pig populations are highly structured and have a small *Ne*. Therefore, increasing only the sample size might not help improve the performance of both variant selection and genomic prediction. Jang et al. [16] reported that using animals with greater EBV reliability (more data available) helped better identify the causative variants than using animals that had lower EBV reliability. Therefore, selecting high-reliability animals and using them could be a possible strategy. Another possible reason for the limited benefit could be the imputation accuracy [19, 38]. The ideal situation to use WGS data is to sequence all the animals in the population without imputation from genotype to sequence level. However, as sequencing the entire population is still impossible, imputation is an inevitable procedure for dealing with WGS data. Since only a limited number of sequenced animals are typically used as a reference for imputation, sequencing more animals and using robust statistical tools to impute alleles accurately are required.

### Comparison of weighted and non-weighted ssGBLUP

WssGBLUP was investigated in addition to ssGBLUP. A major assumption of GBLUP-based methods is that all markers have homogeneous variance. Those methods have been extensively applied for most of the traits in farm animals due to their highly polygenic nature [24, 30]. However, that assumption does not biologically hold because not all markers in the genome explain the same proportion of variance [39]. Therefore, assigning heterogeneous variance per marker for genomic prediction has been investigated in several studies [14, 40, 41]. Weighting SNP in ssGBLUP was initially proposed by Wang et al. [41] by assigning unequal SNP variance through squared SNP effects weighted by allele frequencies. However, this method caused a reduction in GEBV accuracy and extra bias over iteration due to the extreme values of SNP variance, especially for the polygenic traits [42, 43].

Following the increased accuracy reported by Gualdrón-Duarte et al. [14], we used the posterior SNP variances from BayesR as SNP weights. In BayesR, SNP effects are sampled from a mixture of four normal distributions with mean zero and variances equivalent to the following classes: 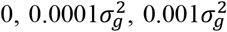 and 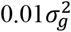 [21]. Therefore, we assumed that this strategy would construct a better weighting matrix close to the true variance of SNP. Our results showed that WssGBLUP outperformed ssGBLUP for ADG, BF, and LDP in TL4 for both Top40k and ChipPlusSign up to 0.06. However, other traits in ML1, ML2, and TL1 showed similar results as in ssGBLUP. Gualdrón-Duarte et al. [14] compared the performances of weighting strategies in Belgian Blue beef cattle. In their study, the average reliability of genomic prediction for 14 traits using GBLUP and BayesR weighted GBLUP showed no differences. However, applying posterior variance of marker effect from the Bayesian mixture model (similar to BayesR) as the weighting factor showed the best performance among other weighting strategies and regular GBLUP in the Nordic Holstein population [44].

The current study showed absent or modest gains in prediction accuracy depending on the lines and traits. We expected almost no gain with WssGBLUP, especially for the largest genotyped population (TL4), as the SNP effects were likely dominated by a large amount of data in the single-step system. However, we observed potential room for improvements in predictions when using the posterior variance of BayesR even with a large amount of data. In other words, although the volume of data could overwhelm *a priori* assumption of SNP effects, improvements can still occur if the variances used as SNP weights are accurate enough.

## Conclusion

Preselection of significant variants from whole-genome sequence data and their utilization could help to improve genomic predictions in maternal and terminal pig lines with tens of thousands of sequenced/imputed animals. However, a limited gain is expected even in large populations. Improvements may be observed when selected variants for some traits are not already represented by the SNP present in commercial SNP chips. Traits with limited accuracy may experience extra gains, given the previous statement holds. Weighting SNP using BayesR variances slightly boosts prediction accuracies. The results of genomic prediction using several preselected variants sets highly depend on the population structure, number of genotyped animals, and method to select variants.

## List of abbreviations

A: pedigree relationship matrix
ADFI: average daily feed intake
ADG: average daily gain
ADGX: ADG recorded in crossbred animals
APY: algorithm for proven and young
BF: backfat thickness
BFX: BF recorded in crossbred animals
dEBV: deregressed EBV
EBV: estimated breeding value
G: genomic relationship matrix
GEBV: genomic EBV
GWAS: genome-wide association study
H: realized relationship matrix
LD: linkage disequilibrium
LDP: loin depth
LDPX: LDP recorded in crossbred animals
ML1: maternal line1
ML2: maternal line2
Ne: effective population size
NSB: number stillborn
QTN: quantitative trait nucleotide
RET: return to oestrus 7 days after weaning
ssGBLUP: single-step GBLUP
SNP: single nucleotide polymorphisms
TL1: terminal line1
TL2: terminal line2
TL3: terminal line3
TL4: terminal line4
TNB: total number of piglets born
WGS: whole-genome sequence
WssGBLUP: weighted-ssGBLUP
WWT: litter weaning weight

## Declarations

### Ethics approval and consent to participate

Not applicable.

### Consent for publication

Not applicable.

### Availability of data and materials

The datasets generated and analysed in this study are from the Pig Improving Company, Genus plc, breeding programme and not publicly available.

## Author’s information

### Contributions

SJ and DL designed the study. RRF and JMH designed and performed the imputation and variant selection steps. CYC assisted in preparing the datasets. SJ performed the analyses and wrote the first draft. DL, IM, RRF, CYC, and WHO provided comments on the manuscript. All authors read and approved the final manuscript.

