## Supplementary file 1 for "Using large-scale whole-genome sequence data for single-step genomic predictions in maternal and terminal pig lines"

**Supplementary table 1. Combination of selected variants for each model**

| Model | SNP panel | Maternal lines | Terminal lines |
|---|---|---|---|
| ADFI | Top40k, ChipPlusSign | ADG + ADFI | ADG + ADFI |
| GROWTH | Top40k, ChipPlusSign | ADG + BFP | ADG + BFP |
| LOINDEPT | Top40k, ChipPlusSign | ADG + LDP | ADG + LDP |
| REPROD | Top40k, ChipPlusSign | TNB + NSB | - |
| RET | Top40k, ChipPlusSign | RET | - |
| WWT | Top40k, ChipPlusSign | WWT | - |

**Supplementary table 2. Number of animals and SNPs for pre-selected SNP panels in maternal lines**

| Model | SNP panel | ML1 | | ML2 | |
|---|---|---|---|---|---|
| | | SNPs | Animals | SNPs | Animals |
| ADFI | ChipPlusSign | 41,364 | 76,227 | 43,325 | 66,608 |
| | Top40k | 80,308 | 76,244 | 80,819 | 66,608 |
| GROWTH | ChipPlusSign | 41,909 | 76,227 | 43,819 | 66,608 |
| | Top40k | 80,613 | 76,230 | 79,984 | 66,608 |
| LOIN | ChipPlusSign | 41,742 | 76,227 | 43,618 | 66,608 |
| | Top40k | 79,605 | 76,230 | 79,625 | 66,608 |
| REPROD | ChipPlusSign | 40,707 | 76,227 | 42,907 | 66,608 |
| | Top40k | 41,180 | 76,214 | 41,173 | 66,561 |
| RET | ChipPlusSign | 40,624 | 76,227 | 42,819 | 66,608 |
| | Top40k | 41,171 | 76,177 | 41,168 | 66,300 |
| WWT | ChipPlusSign | 40,772 | 76,227 | 42,777 | 66,608 |
| | Top40k | 41,173 | 76,244 | 41,165 | 66,608 |

**Supplementary table 3. Number of animals and SNPs for pre-selected SNP panels in terminal lines**

| Model | SNP panel | TL1 | | TL2 | | TL3 | | TL4 | |
|---|---|---|---|---|---|---|---|---|---|
| | | **SNPs** | **Animals** | **SNPs** | **Animals** | **SNPs** | **Animals** | **SNPs** | **Animals** |
| ADFI | ChipPlusSign | 36,305 | 60,467 | 40,744 | 41,572 | 40,267 | 29,328 | 43,814 | 104,644 |
| | Top40k | 79,594 | 59,453 | 78,714 | 41,507 | 80,070 | 29,195 | 80,153 | 104,659 |
| GROWTH | ChipPlusSign | 36,792 | 60,467 | 41,578 | 41,572 | 40,432 | 29,328 | 44,734 | 104,645 |
| | Top40k | 79,078 | 59,593 | 79,112 | 41,533 | 80,155 | 29,307 | 80,899 | 104,659 |
| LOIN | ChipPlusSign | 36,796 | 60,467 | 40,798 | 41,572 | 40,429 | 29,328 | 44,305 | 104,645 |
| | Top40k | 78,789 | 59,615 | 77,878 | 41,535 | 80,535 | 29,286 | 81,439 | 104,659 |

**Supplementary table 4. Number of test animals for each trait in all lines**

| Line | ADFI | ADG | BF | LDP | TNB | NSB | RET | WWT |
|---|---|---|---|---|---|---|---|---|
| ML1 | 8,387 | 10,614 | 8,418 | 8,422 | 425 | 362 | 246 | 332 |
| ML2 | 6,976 | 9,237 | 9,237 | 7,363 | 399 | 401 | 220 | 282 |
| **Line** | **ADFI** | **ADG** | **BF** | **ADGX** | **BFX** | **LDP** | | **LDPX** |
| TL1 | 5,970 | 5,970 | 5,970 | 5,970 | 5,970 | 5,965 | | 5,943 |
| TL2 | 3,720 | 3,720 | 3,720 | 3,720 | 3,720 | 3,720 | | 2,858 |
| TL3 | 1,808 | 2,324 | 2,324 | 2,324 | 2,324 | 2,324 | | 2,324 |
| TL4 | 9,434 | 11,308 | 11,308 | 11,308 | 11,308 | 11,308 | | 11,308 |

**Supplementary table 5. Prediction accuracy of maternal lines using ssGBLUP**

| Line | SNP panel | ADFI | ADG | BF | LDP | TNB | NSB | RET | WWT |
|---|---|---|---|---|---|---|---|---|---|
| ML1 | Chip | 0.361 | 0.460 | 0.509 | 0.515 | 0.406 | 0.358 | 0.139 | 0.304 |
| | ChipPlusSign | 0.366 | 0.468 | 0.514 | 0.519 | 0.410 | 0.360 | 0.139 | 0.303 |
| | Top40k | 0.368 | 0.488 | 0.515 | 0.530 | 0.419 | 0.358 | 0.187 | 0.288 |
| ML2 | Chip | 0.371 | 0.611 | 0.629 | 0.611 | 0.348 | 0.366 | 0.204 | 0.284 |
| | ChipPlusSign | 0.374 | 0.609 | 0.634 | 0.620 | 0.351 | 0.371 | 0.204 | 0.282 |
| | Top40k | 0.355 | 0.608 | 0.644 | 0.618 | 0.368 | 0.392 | 0.251 | 0.285 |

**Supplementary table 6. Accuracy gain and reduction (%) compared to Chip data in maternal lines**

| Line | SNP panel | ADFI | ADG | BF | LDP | TNB | NSB | RET | WWT | Mean |
|---|---|---|---|---|---|---|---|---|---|---|
| ML1 | ChipPlusSign | 1.34 | 1.61 | 0.91 | 0.66 | 1.06 | 0.56 | 0.12 | -0.26 | 0.75 |
| | Top40k | 1.73 | 5.94 | 1.12 | 2.78 | 3.14 | 0.04 | 34.77 | -5.19 | 5.54 |
| ML2 | ChipPlusSign | 0.78 | -0.38 | 0.74 | 1.44 | 0.99 | 1.49 | -0.40 | -0.74 | 0.49 |
| | Top40k | -4.25 | -0.56 | 2.39 | 1.07 | 5.62 | 7.21 | 22.87 | 0.34 | 4.34 |

**Supplementary table 7. Prediction accuracy of terminal lines using ssGBLUP**

| Line | SNP panel | ADFI | ADG | BF | ADGX | BFX | LDP | LDPX |
|---|---|---|---|---|---|---|---|---|
| TL1 | Chip | 0.359 | 0.488 | 0.600 | 0.334 | 0.616 | 0.596 | 0.321 |
| | ChipPlusSign | 0.363 | 0.494 | 0.608 | 0.337 | 0.618 | 0.600 | 0.324 |
| | Top40k | 0.338 | 0.427 | 0.563 | 0.308 | 0.625 | 0.548 | 0.304 |
| TL2 | Chip | 0.295 | 0.483 | 0.600 | 0.178 | 0.481 | 0.473 | 0.050 |
| | ChipPlusSign | 0.301 | 0.492 | 0.607 | 0.182 | 0.489 | 0.474 | 0.047 |
| | Top40k | 0.301 | 0.504 | 0.598 | 0.207 | 0.474 | 0.465 | 0.029 |
| TL3 | Chip | 0.356 | 0.629 | 0.546 | 0.392 | 0.414 | 0.611 | 0.572 |
| | ChipPlusSign | 0.360 | 0.632 | 0.547 | 0.394 | 0.416 | 0.611 | 0.570 |
| | Top40k | 0.366 | 0.640 | 0.548 | 0.401 | 0.427 | 0.634 | 0.585 |
| TL4 | Chip | 0.397 | 0.497 | 0.594 | 0.262 | 0.587 | 0.564 | 0.483 |
| | ChipPlusSign | 0.403 | 0.505 | 0.599 | 0.269 | 0.592 | 0.571 | 0.491 |
| | Top40k | 0.395 | 0.510 | 0.601 | 0.283 | 0.593 | 0.587 | 0.517 |

**Supplementary table 8. Accuracy gain and reduction (%) compared to Chip data in terminal lines**

| Line | SNP panel | ADFI | ADG | BF | ADGX | BFX | LDP | LDPX | Mean |
|---|---|---|---|---|---|---|---|---|---|
| TL1 | ChipPlusSign | 1.04 | 1.30 | 1.32 | 0.95 | 0.31 | 0.71 | 1.05 | 0.95 |
| | Top40k | -5.80 | -12.49 | -6.16 | -7.74 | 1.55 | -8.05 | -5.28 | -6.28 |
| TL2 | ChipPlusSign | 2.20 | 1.85 | 1.18 | 2.27 | 1.53 | 0.14 | -5.26 | 0.56 |
| | Top40k | 2.20 | 4.36 | -0.29 | 16.16 | -1.51 | -1.63 | -42.56 | -3.32 |
| TL3 | ChipPlusSign | 0.94 | 0.47 | 0.29 | 0.66 | 0.69 | 0.00 | -0.45 | 0.37 |
| | Top40k | 2.74 | 1.74 | 0.44 | 2.35 | 3.18 | 3.74 | 2.24 | 2.35 |
| TL4 | ChipPlusSign | 1.65 | 1.69 | 0.96 | 2.61 | 0.76 | 1.22 | 1.63 | 1.50 |
| | Top40k | -0.49 | 2.54 | 1.20 | 7.91 | 0.91 | 3.95 | 7.07 | 3.30 |

**Supplementary table 9. $b_1$ of maternal lines using ssGBLUP**

| Line | SNP panel | ADFI | ADG | BF | LDP | TNB | NSB | RET | WWT | Mean |
|---|---|---|---|---|---|---|---|---|---|---|
| ML1 | Chip | 0.91 | 0.56 | 0.61 | 0.66 | 0.80 | 0.75 | 0.47 | 0.71 | 0.68 |
| | ChipPlusSign | 0.90 | 0.56 | 0.61 | 0.66 | 0.80 | 0.76 | 0.47 | 0.70 | 0.68 |
| | Top40k | 0.89 | 0.60 | 0.64 | 0.68 | 0.72 | 0.85 | 0.47 | 0.65 | 0.69 |
| ML2 | Chip | 0.99 | 0.77 | 0.67 | 0.79 | 0.77 | 1.06 | 0.56 | 0.80 | 0.80 |
| | ChipPlusSign | 0.98 | 0.74 | 0.64 | 0.77 | 0.78 | 1.07 | 0.56 | 0.79 | 0.79 |
| | Top40k | 0.90 | 0.73 | 0.69 | 0.77 | 0.89 | 1.23 | 0.68 | 0.72 | 0.83 |

**Supplementary table 10. $b_1$ of terminal lines using ssGBLUP**

| Line | SNP panel | ADFI | ADG | BF | ADGX | BFX | LDP | LDPX | Mean |
|---|---|---|---|---|---|---|---|---|---|
| TL1 | Chip | 0.74 | 0.62 | 0.70 | 0.76 | 0.58 | 0.69 | 0.50 | 0.66 |
| | ChipPlusSign | 0.74 | 0.63 | 0.70 | 0.76 | 0.59 | 0.70 | 0.51 | 0.66 |
| | Top40k | 0.68 | 0.64 | 0.72 | 0.77 | 0.79 | 0.69 | 0.67 | 0.71 |
| TL2 | Chip | 0.63 | 0.64 | 0.58 | 0.79 | 0.60 | 0.60 | 1.23 | 0.72 |
| | ChipPlusSign | 0.64 | 0.64 | 0.57 | 0.80 | 0.60 | 0.60 | 1.16 | 0.72 |
| | Top40k | 0.57 | 0.59 | 0.57 | 0.85 | 0.58 | 0.55 | 0.43 | 0.59 |
| TL3 | Chip | 1.08 | 0.68 | 0.63 | 0.91 | 0.68 | 0.60 | 0.66 | 0.75 |
| | ChipPlusSign | 1.08 | 0.68 | 0.63 | 0.91 | 0.68 | 0.59 | 0.69 | 0.75 |
| | Top40k | 1.07 | 0.66 | 0.62 | 0.95 | 0.73 | 0.55 | 0.65 | 0.75 |
| TL4 | Chip | 0.86 | 0.65 | 0.65 | 0.62 | 0.68 | 0.71 | 0.66 | 0.69 |
| | ChipPlusSign | 0.85 | 0.65 | 0.64 | 0.63 | 0.67 | 0.71 | 0.67 | 0.69 |
| | Top40k | 0.80 | 0.66 | 0.66 | 0.75 | 0.72 | 0.71 | 0.74 | 0.72 |

**Supplementary table 11. $b_1$ of WssGBLUP compared to ssGBLUP**

| Lines | Description | ADFI | ADG | BF | LDP |
|---|---|---|---|---|---|
| ML1 | Top40k | 0.89 | 0.60 | 0.64 | 0.68 |
| | Top40k weighted | 0.89 | 0.59 | 0.64 | 0.68 |
| | ChipPlusSign | 0.90 | 0.56 | 0.61 | 0.66 |
| | ChipPlusSign weighted | 0.98 | 0.56 | 0.62 | 0.66 |
| ML2 | Top40k | 0.90 | 0.73 | 0.69 | 0.77 |
| | Top40k weighted | 0.88 | 0.72 | 0.66 | 1.10 |
| | ChipPlusSign | 0.98 | 0.74 | 0.64 | 0.77 |
| | ChipPlusSign weighted | 0.99 | 0.74 | 0.63 | 1.10 |
| TL1 | Top40k | 0.68 | 0.64 | 0.72 | 0.69 |
| | Top40k weighted | 0.69 | 0.64 | 0.72 | 0.69 |
| | ChipPlusSign | 0.74 | 0.63 | 0.70 | 0.70 |
| | ChipPlusSign weighted | 0.74 | 0.62 | 0.71 | 0.70 |
| TL4 | Top40k | 0.86 | 0.65 | 0.65 | 0.71 |
| | Top40k weighted | 0.79 | 0.65 | 0.65 | 0.71 |
| | ChipPlusSign | 0.80 | 0.66 | 0.64 | 0.71 |
| | ChipPlusSign weighted | 0.84 | 0.64 | 0.63 | 0.70 |
